# Stochastic LASSO for extremely high-dimensional genomic data

**DOI:** 10.1101/2025.04.22.650135

**Authors:** Beomsu Baek, Jongkwon Jo, Mingon Kang, Youngsoon Kim

**Affiliations:** Department of Information and Statistics, Gyeongsang National University, Jinju, Republic of Korea; Department of Computer Science, University of Nevada, Las Vegas, Las Vegas, NV 89154, USA

**Keywords:** Stochastic LASSO, LASSO, high-dimensional data, variable selection

## Abstract

Accurate identification of significant features in high-dimensional data is indispensable in high-throughput genomic analysis and association studies. Least Absolute Shrinkage and Selection Operator (LASSO) and its derivatives have been widely adapted to discover potential biomarkers as a feature selection scheme in various biological systems. Recently, bootstrap-based LASSO models, such as Random LASSO and Hi-LASSO, have been effective solutions for extremely high-dimensional but low sample size (EHDLSS) genomic data. However, the bootstrap-based LASSO models still have several drawbacks, such as multicollinearity within bootstrap samples, missing predictors in draw, and randomness in predictor sampling. To tackle the limitations, we propose a new bootstrap-based LASSO, named Stochastic LASSO, that effectively reduces multicollinearity in bootstrap samples and mitigates randomness in predictor sampling, resulting in remarkably outperforming benchmarks in feature selection and coefficient estimation. Furthermore, Stochastic LASSO provides a two-stage t-test strategy for selecting statistically significant features. The performance of Stochastic LASSO was assessed by comparing the existing benchmark models in extensive simulation experiments. In the simulation experiments, Stochastic LASSO consistently showed significant improvements in performance compared to the state-of-the-art LASSO models for feature selection, coefficient estimation, and robustness. We also applied Stochastic LASSO for the gene expression data of publicly available TCGA cancer datasets and identified statistically significant genes associated with survival month prediction. The source code is publicly available at: https://github.com/datax-lab/StochasticLASSO.

## 1 Introduction

Identification of a subset of significant predictors is indispensable in understanding biological mechanisms and enhancing predictive performance in high-throughput and high-dimensional genomic data [1–3]. Least Absolute Shrinkage and Selection Operator (LASSO) [4] and its derivatives have powered to identify a subset of relevant predictors in several biological applications, such as survival analysis [5], metabolomics [6], discovering protein–protein interactions [7], and Cox proportional hazard modeling [8]. However, conventional LASSO models, including Elastic-Net [9], Adaptive LASSO [10], Relaxed LASSO [11], and Precision LASSO [12], have following limitations when applied to extremely high-dimensional and low sample size (EHDLSS) data: (1) LASSO selects predictors only up to the sample size, and (2) LASSO does not identify all the predictors that are highly correlated with each other. Thus, the conventional LASSOs are challenged to apply for omics datasets that include hundreds of patent samples of more than 20,000 genes or 80,000 SNPs with highly multicollinearity.

Bootstrap-based LASSOs, such as Random LASSO [13], Recursive Random LASSO [14], and Hi-LASSO [15], addressed the EHDLSS issues by drawing multiple bootstrap samples of lower-dimensionality and then aggregating the results for the final feature selection. Most bootstrap-based LASSOs consist of two procedures: (1) calculating importance scores of predictors for the oracle property [16] and (2) estimating coefficients of predictors by prioritizing predictors of high importance scores. The bootstrap-based LASSOs have the advantages of accurate feature selection, precise coefficient estimation, and enhanced predictive performance with EHDLSS data.

However, the bootstrap-based LASSOs also have several drawbacks, such as a multicollinearity issue within bootstrap samples, missing predictors in draw, and randomness in predictor sampling. First, multicollinearity within bootstrap samples often causes underestimated importance scores of non-zero predictors. Although bootstrap-based LASSOs reduce multicollinearity in the entire set of predictors through bootstrap sampling, the bootstrap samples still have local multicollinearity. For instance, non-zero predictors are often estimated as zero, when highly correlated non-zero predictors are drawn in a bootstrap sample. Moreover, if highly correlated predictors are with opposite coefficient signs, only the predictors with dominant sign are identified. The predictors of a non-dominant sign are estimated as zero or incorrectly identified as the dominant sign. Second, bootstrap-based LASSOs have a potential risk of missing predictors during the bootstrap process. Let the model construct *B* numbers of bootstrap samples by drawing *q* predictors from a total of *p* predictors (*q < p*). The number of times that each predictor is drawn follows the binomial distribution. Then, when 500 bootstrap samples are generated by drawing 200 predictors from 20,000 predictors (e.g., *B* = 500, *q* = 200, and *p* = 20, 000), the expected number of predictors that are never selected is 131.4, and the expected number of predictors selected less than three times is 2,467.7. Thus, a number of the predictors’ coefficients would be missing or underestimated, since the majority of predictors are seldom drawn. Lastly, missing predictors or imbalanced predictor inclusion in the bootstrap, caused by randomness in predictor sampling, make coefficient estimations ineffective. Bootstrap-based LASSOs construct bootstrap samples by drawing *q* predictors with probabilities proportional to the importance scores. However, the randomness in draw still makes chances of several non-zero predictors missing or seldom consideration from the regression model in a bootstrap.

In this paper, we propose an enhanced bootstrap-based LASSO model, named Stochastic LASSO, which remarkably improves the current LASSO solutions. The main contributions of Stochastic LASSO are as follows:

- Stochastic LASSO significantly enhances the feature selection performance, as well as accurately estimating true coefficients, comparing to the state-of-the-art LASSO models,
- Stochastic LASSO proposes a parametric statistical test for selecting significant feature in high-dimensional data, and
- Stochastic LASSO produces robust feature selection results.

The rest of the paper is organized as follows: Section 2 describes the proposed Stochastic LASSO in detail, and in Section 3, we conducted the assessment of Stochastic LASSO by comparing it with existing state-of-the-art LASSO models.

## 2 Methods

### 2.1 2.1. Overview

Stochastic LASSO follows the procedures: (1) constructing lower-dimensional, linearly independent bootstrap samples by a Correlation Based Bootstrapping (CBB) strategy, (2) estimating coefficients of each bootstrap sample, (3) calculating local scores for feature selection, (4) estimating the final coefficients (i.e., global scores) by forward selection, and (5) determining statistical significance by a two-stage t-test in a one-time bootstrapping procedure. Stochastic LASSO improves the LASSO solution by proposing (1) an enhanced bootstrapping algorithm for reducing multicollinearity (§2.2), (2) a forward selection strategy for robust feature selection and coefficient estimation (§2.3), and (3) a statistical strategy for identifying statistically significant features in high dimensional data (§2.4).

### 2.2 2.2. Reducing multicollinearity in bootstrap samples

Stochastic LASSO effectively reduces multicollinearity in bootstrap samples by the proposed Correlation Based Bootstrapping (CBB) algorithm. CBB penalizes predictors highly correlated with others in the bootstrapping so that the predictors of bootstrap samples become independent. CBB sets a selection probability of each predictor by the correlation with other predictors in the bootstrap sample, when Stochastic LASSO draws a predictor during bootstrap sampling. Let *S* and *Q* be the sets of the predictor indices, where *S* includes predictor indices that have not been drawn and *Q* is with indices already included in the bootstrap sample. Initially, *S* = {1, …, *p*} and *Q* = ∅. Then, CBB computes the selection probabilities of *S* based on correlation with *Q*. We refine the probability *Pr*(*p*_*i*_) (where *i* ∈ *S*) that the *i*-th predictor is selected, as follows:

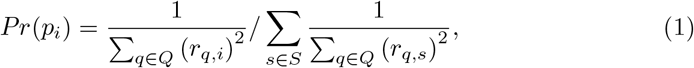

where *r*_*i,j*_ denotes the Pearson correlation coefficient between *i*-th and *j*-th predictors. Once an *i*-th predictor is randomly selected with *Pr*(*p*_*i*_), CBB updates *S* and *Q* as *S* = *S*\{*i*} and *Q* = *Q* ∪ *i* until the bootstrap sample is constructed with *q* predictors (i.e., |*Q*| = *q*), where *S*\{*i*} is the vector *S* excluding the element *i*. When Stochastic LASSO draws the first predictor of the bootstrap sample (i.e., *Q* = ∅), CBB sets *Pr*(*p*_*i*_) to 1*/* |*S*|. Therefore, CBB prioritizes predictors that are likely independent to the other predictors in the bootstrapping.

In addition, Stochastic LASSO guarantees that all predictors are drawn equal times by sampling predictors without replacement. CBB algorithm constructs ⌈*p/q*⌉ boot-strap samples until *S* = ∅, ensuring that each predictor is drawn once. To guarantee that each predictor is drawn exactly same times, Stochastic LASSO repeats the CBB algorithm *r* times. Consequently, Stochastic LASSO estimates *r* coefficients for each of the *p* predictors by applying penalized linear regression to each bootstrap samples.

### 2.3 2.3. Improving coefficient estimation

Stochastic LASSO determines the optimal subset of features and estimates their coefficients using forward selection based on local scores. While traditional forward selection methods require evaluating all possible feature subsets, Stochastic LASSO defines a fixed subset based on the rank of local scores. Let 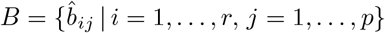 be a *r*×*p* matrix that includes *r* coefficient estimates of *p* numbers of variables obtained from bootstrapping procedure. We define the local score of the *j*-th predictor (*L*_*j*_) as follows:

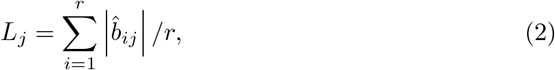

Let *S*_0_ = ∅ be an initial feature subset, and the subsequent subsets (*S*_*k*_) are defined by prioritizing features with high local scores:

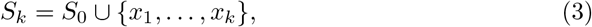

where *x*_*j*_ denotes the predictor with the *j*-th highest local score. Then, Stochastic LASSO estimates the coefficients of the feature subsets (*S*_*k*_) by applying penalized linear regression (e.g., Elastic-Net), and determines the optimal subset *S*^+^ as follows:

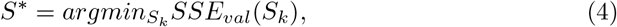

where *SSE*_*val*_(*S*_*k*_) denotes the sum of squared error of validation data calculated by the estimated coefficients of *S*_*k*_. Thus, Stochastic LASSO systematically constructs a subset consisting of non-zero predictors and precisely estimates their partial correlations, without additional bootstrapping procedures that most bootstrap-based LASSOs require.

### 2.4 2.4. Tests of significance for statistically significant feature

Stochastic LASSO proposes a statistical test, two-stage t-test (TSTT), for evaluating statistical significance of features. TSTT sequentially conducts one-sample t-test and two-sample t-test. The one-sample t-test identifies potential significant features whose estimated coefficient means are non-zeros, and the two-sample t-test selects significant features with a large coefficient scale among the potential significant features. Let *β*_*j*_ be a regression coefficient of the *j*-th feature. The coefficient estimate of the *j*-th feature, 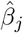, is defined as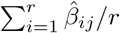, where 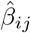 denotes the *i*-th coefficient estimate of the *j*-th feature. Then the null and alternative hypotheses for the one-sample test are as follows:

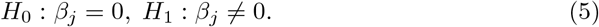

Let *K* be a set of indices of the *m* numbers of potential significant features selected through the previous one-sample t-test. The estimate of absolute value of coefficient of *k*-th feature, 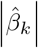 is defined as 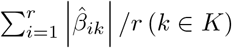. The estimate of absolute value of coefficient of all potential significant features, 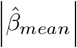, is defined as 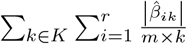.Then the null and alternative hypotheses for the two-sample t-test are as follows:

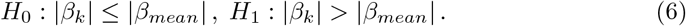

The detail procedures of Stochastic LASSO are described in Algorithm 1.

#### Algorithm 1 Stochastic LASSO

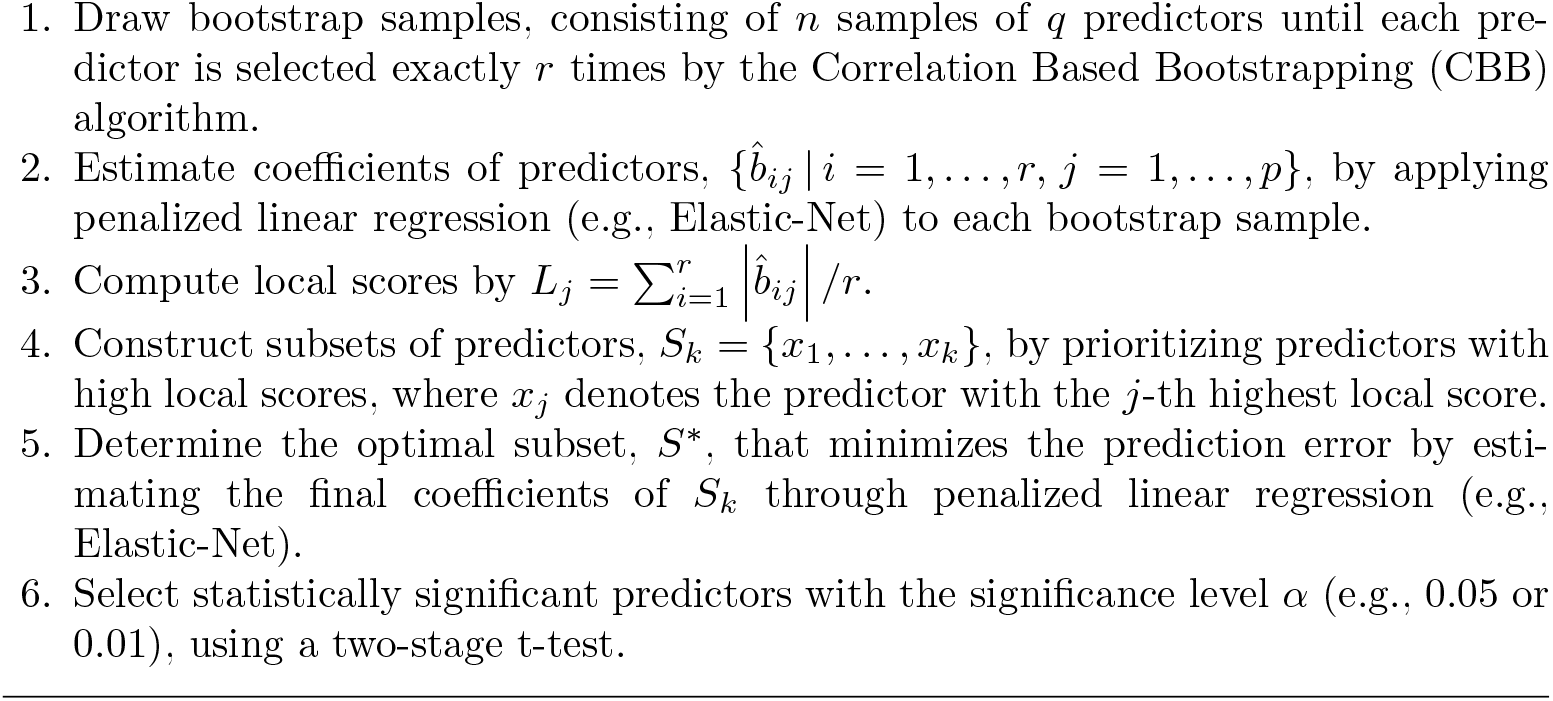

## 3 Experimental Results

We assessed the performance of Stochastic LASSO with the various experimental settings, compared to state-of-the-art LASSO models on the following criteria: (1) feature selection, (2) coefficient estimation, (3) test of significance, and (4) model robustness. First, we evaluated how accurately Stochastic LASSO identifies non-zero variables with various scales of data dimensionalities. Second, we assessed the performance of coefficient estimation by computing the residual sum between the estimations and ground truth. Third, we examined whether Stochastic LASSO can identify statistical significance of non-zero variables. Lastly, we evaluated the consistency and robustness of feature selection.

### 3.1 3.1. Feature selection

For the assessment of feature selection, we conducted simulation studies where ground truth of non-zero variables are known. The simulation study mainly considered high-dimensional, but low-sample-size data with high multicollinearity, which is a common setting in genomic data. We followed the simulation settings that have been commonly used in most bootstrap-based LASSO studies [13–15]. We generated four synthetic datasets (i.e., Dataset I-IV) with the following linear regression model:

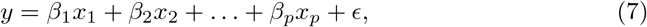

where *ϵ* ∼ *N* (0, *σ*^2^), *x*_*i*_ ∼ *N*(0, 1), and various types of multicollinearity were introduced by predefined covariance matrices. The four dataset consisted of varying numbers of variables and samples, defined in Table 1. Dataset I included 50 samples of 100 variables, where the first ten coefficients were set to non-zeros. The regression coefficients of ground truth were defined as:

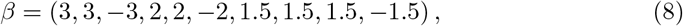

and the multicollinearity was introduced by predefined covariance matrix:

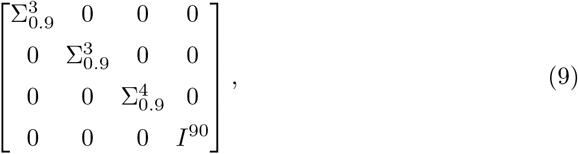

where 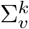 is a *k* × *k* matrix with unit diagonal elements and off-diagonal elements of value *v*, and *I*^*k*^ is an identity matrix of size *k*. Dataset II consists of 100 samples of 1,000 variables, where the first 50 non-zero coefficients were drawn from *N* (0, 4). The multicollinearity with high and low degrees was designed using the following covariance matrix:

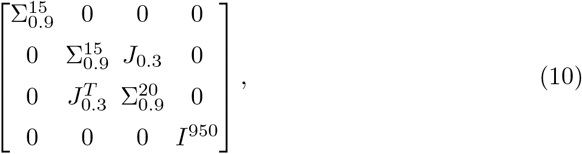

where *J*_*v*_ is a matrix with all unit elements of a value *v*. Dataset III includes 200 samples of 10,000 variables, where the first 50 coefficients of non-zero were drawn from *N* (0, 4). The covariance matrix of Dataset III is as follows:

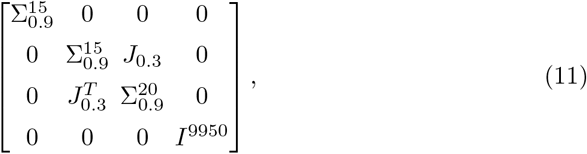

**Table 1.** Description of the simulation data.

| 2-34-5 | Sample sizes | | Numbers of features | | $\beta$ |
| --- | --- | --- | --- | --- | --- |
|  | Training | Validation* | Total | Non-zeros |  |
| Dataset I | 50 | 10 | 100 | 10 | $(3, 3, -3, 2, 2, -2, 1.5, 1.5, 1.5, -1.5, 0, \dots, 0)$ |
| Dataset II | 100 | 20 | 1,000 | 50 | $\beta_1, \dots, \beta_{50} \sim N(0, 4)$, others zero |
| Dataset III | 200 | 40 | 10,000 | 50 | $\beta_1, \dots, \beta_{50} \sim N(0, 4)$, others zero |
| Dataset IV | 400 | 80 | 10,000 | 50 | $\beta_1, \dots, \beta_{50} \sim N(0, 4)$, others zero |

\*The validation data is further generated with the size equivalent to 20% of the training data.

In Dataset IV, we considered double samples in the same model of Dataset III. To tune the hyper-parameters of the LASSO models, we further generated validation data for each dataset, with a size equivalent to 20% of the training samples. Note that test data was not considered, as we evaluated only feature selection performance without assessing predictive errors.

The benchmark models include LASSO [4], Elastic-Net [9], Adaptive LASSO [10], Relaxed LASSO [11], Precision LASSO [12], Random LASSO [13], Recursive Random LASSO [14], and Hi-LASSO [15]. For the non-bootstrap-based LASSO models (i.e., LASSO, Elastic-Net, Adaptive LASSO, Relaxed LASSO, and Precision LASSO), hyper-parameters (e.g., regularization parameter) were tuned by minimizing prediction error on the validation data. For the bootstrap-based LASSO models of Random LASSO, Recursive Random LASSO, and Hi-LASSO, the number of variables (*q*_1_ and *q*_2_) were set to the sample number (i.e., *n*). The number of bootstrap samples (*B*) was set to *p/q* × 30 to ensure that each variable is included in the bootstrap samples 30 times on average. In Stochastic LASSO, we also set *q* = *n* and *r* = 30 for the fair comparison.

We computed F1-scores and Area Under the Precision-Recall Curve (AUCPR) for the evaluation. We defined non-zero variables as positive, and zero variables as negative. Then, the confusion matrices were computed as follows: True Positive (*TP*) if a model correctly identifies non-zero variables as non-zeros; False Positive (*FP*) if a model incorrectly identifies zero variables as non-zeros; False Negative (*FN*) if a model incorrectly identifies non-zero variables as zeros; and True Negative (*TN*) if a model correctly identifies zero variables as zeros. F1-score was calculated by 2(*Precision* × *Recall*)/(*Precision* + *Recall*), where *Precision* and *Recall* are defined as *TP/*(*TP* + *FP*) and *TP/*(*TP* +*FN*), respectively. The AUCPR was calculated from the Precision-Recall curve, which is generated by evaluating the model across varying thresholds. We repeated the experiments ten times on randomly generated synthetic data for the reproducibility of the model performance.

Stochastic LASSO outperformed all other benchmarks across the synthetic datasets (Fig. 1A, Supplementary Table S1), showing the highest F1-scores of 0.7093±0.0197, 0.6650±0.0139, 0.5251±0.0170, and 0.7777±0.0046 with Dataset I-IV, respectively, which showed 8%, 77%, 65%, and 42% improvements against the second-best models. The outperformance of Stochastic LASSO to the second-best benchmark was statistically validated by the Wilcoxon rank-sum test (p-values*<*0.05) with all synthetic datasets (Supplementary Table S1). Furthermore, we verified the feature selection performance using AUCPR without thresholding. Stochastic LASSO also achieved the highest AUCPR of 0.8284±0.0095, 0.6992±0.0021, 0.5772±0.0089, and 0.6985± 0.0018 with Dataset I-IV, respectively, representing statistically significant (p-values*<*0.05) improvements of 15%, 37%, 31%, and 8% over the second-best models (Fig. 1B, Supplementary Table S1). We observed the remarkable performance of bootstrap-based LASSO models, including Random LASSO and Hi-LASSO, in the high dimensional data (e.g., Dataset II-IV), which implies that bootstrap-based LASSO is more suitable for extremely high-dimensional data analysis than non-bootstrap-based LASSO. However, Recursive Random LASSO showed the lowest F1-scores, despite being a bootstrap-based model, due to introducing initial biases when handling high-dimensional data.

**Fig. 1.**
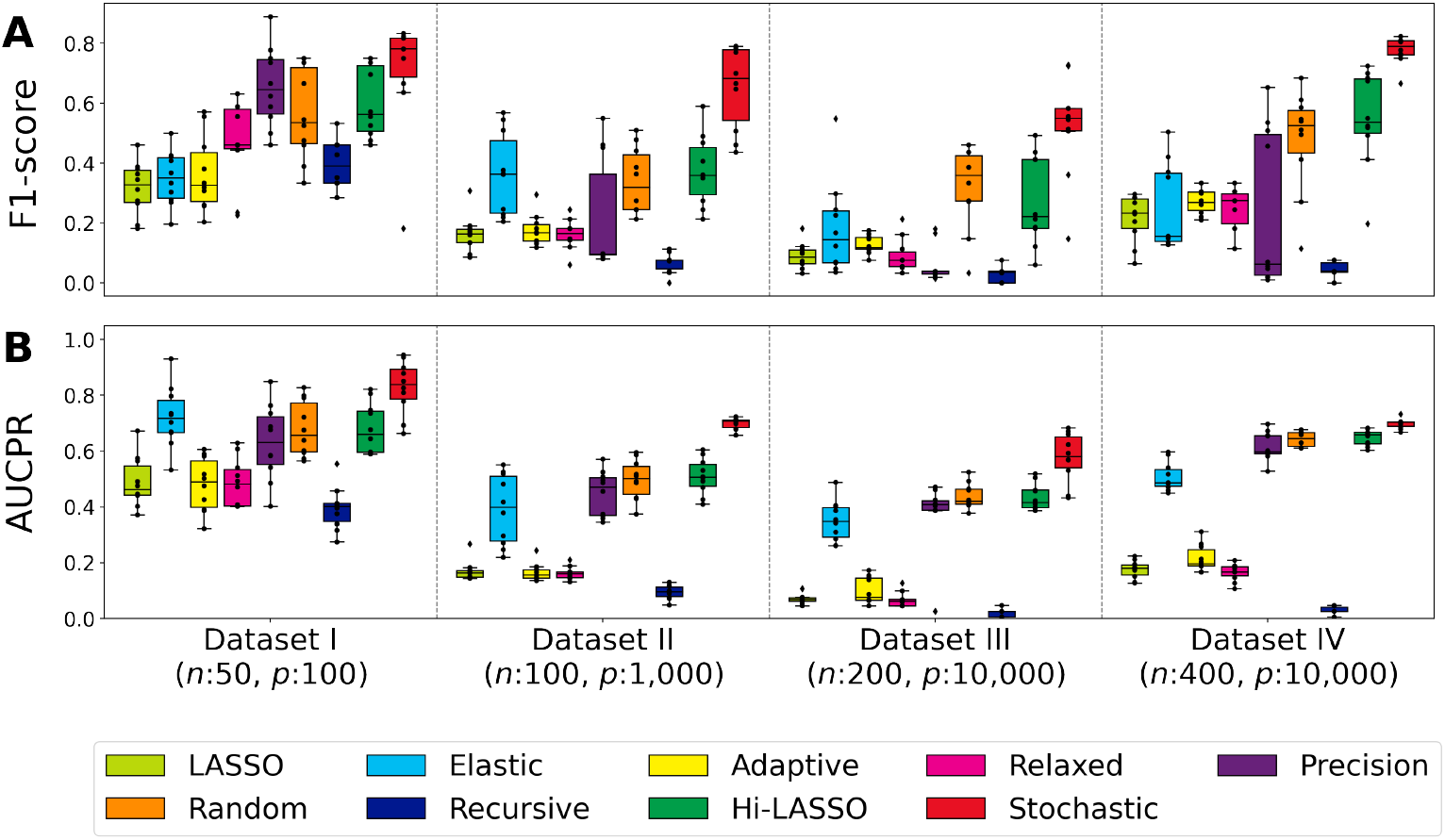
Feature selection performance with simulation data. (A) Comparison of F1-score, (B) Comparison of AUCPR. On the x-axis of the figure, we labeled the sample size (*n*) and dimension (*p*) for each dataset.

### 3.2 3.2. Coefficient estimation

In this experiment, we verified how precisely the models estimate the ground truth coefficient values. The performance of coefficient estimation was evaluated by computing Root Mean Squared Errors (RMSE) between estimated coefficients and the ground truth in the synthetic datasets that we used in the previous experiment (Dataset I-IV). We computed *RMSE*_*ALL*_ and *RMSE*_*Nonzeros*_, where *RMSE*_*ALL*_ was computed with all coefficients, and *RMSE*_*Nonzeros*_ was computed with only non-zero coefficients. Stochastic LASSO showed the least errors to estimate coefficient values among the benchmark models (Fig. 2, Supplementary Table S2), achieving the lowest *RMSE*_*ALL*_ of 0.5022±0.0091, 0.3596±0.0041, 0.1099±0.0006, and 0.0777±0.0002 with Dataset I-IV, respectively, which represented 9%, 5%, 9% and 8% improvements over the second-best models. Stochastic LASSO also exhibited the lowest *RMSE*_*Nonzeros*_ of 1.5286±0.0294, 1.5931±0.0179, 1.5464±0.0098, and 1.0945±0.0025 with Dataset I-IV, respectively, showing 12%, 6%, 9%, and 8% improvements against the second-best models. Stochastic LASSO’s improvements for coefficient estimation were statistically validated compared to the second-best models (p-values*<*0.05). The second-best models were Hi-LASSO in Dataset I&II and Random LASSO in Dataset III&IV.

**Fig. 2.**
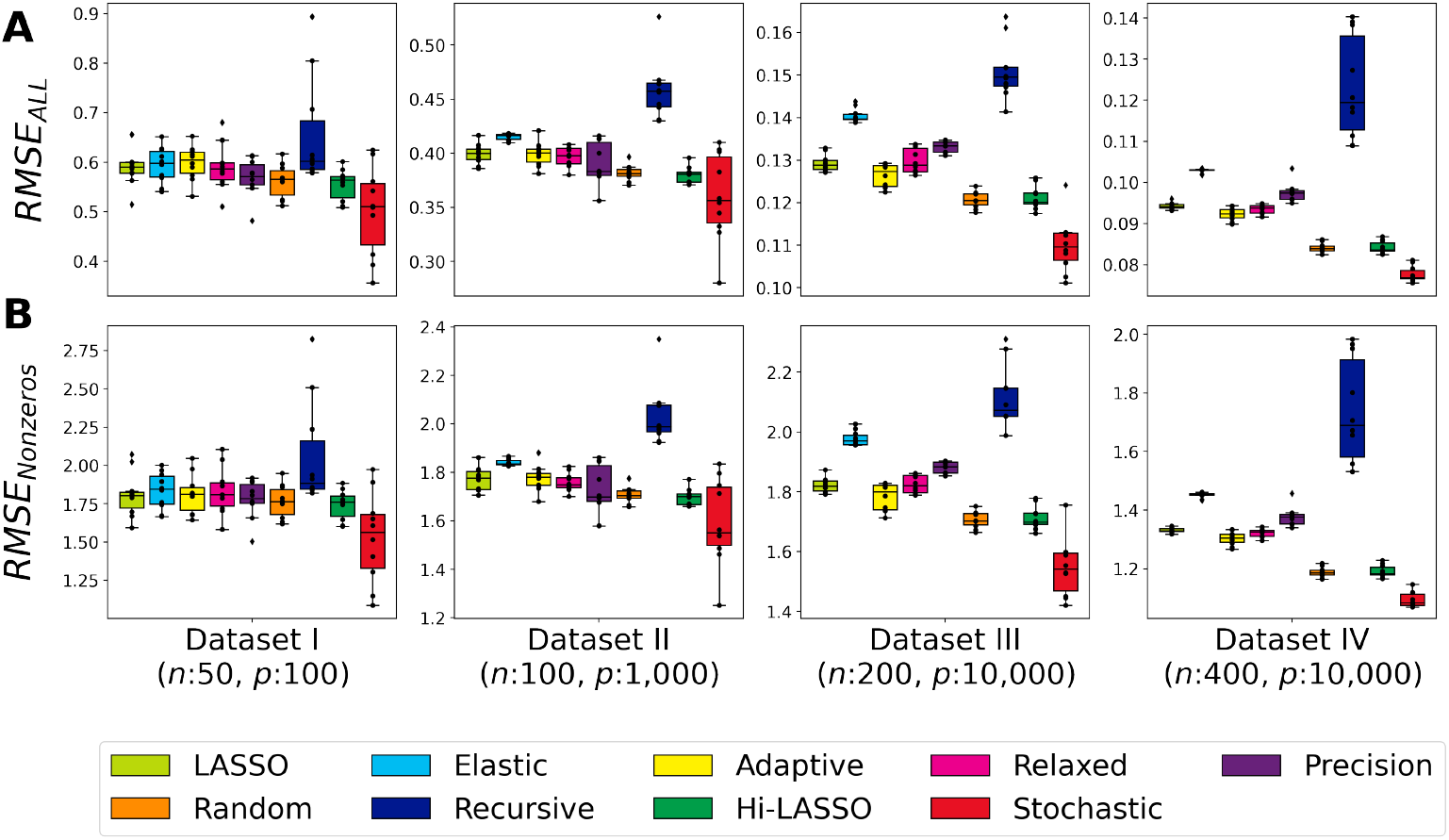
Coefficient estimation performance with simulation data. (A) Comparison of *RMSE*_*ALL*_, (B) Comparison of *RMSE*_*Nonzeros*_. On the x-axis of the figure, we labeled the sample size (*n*) and dimension (*p*) for each dataset.

Furthermore, we explored the signs of Stochastic LASSO’s coefficients compared to the second-best models of Hi-LASSO and Random LASSO. Fig. 3 depicts the results of the coefficient estimations of the models. In Fig. 3, the circle marker in black presents the ground truth of non-zero coefficients, while colored shape markers indicate the estimations of the models on average (i.e., Stochastic LASSO with pentagon in red, Hi-LASSO with square in green, and Random LASSO with triangle in orange). The figure shows that Stochastic LASSO more accurately estimated the signs of the coefficients than the others. In Dataset I, Random LASSO and Hi-LASSO failed to estimate the negative signs on *β*_3_, *β*_6_, and *β*_10_, whereas Stochastic LASSO successfully estimated them (Fig. 3A). Note that the non-zero variables in Dataset I are designed with predominantly positive coefficients and high multicollinearity (Equation 8 & 9).

**Fig. 3.**
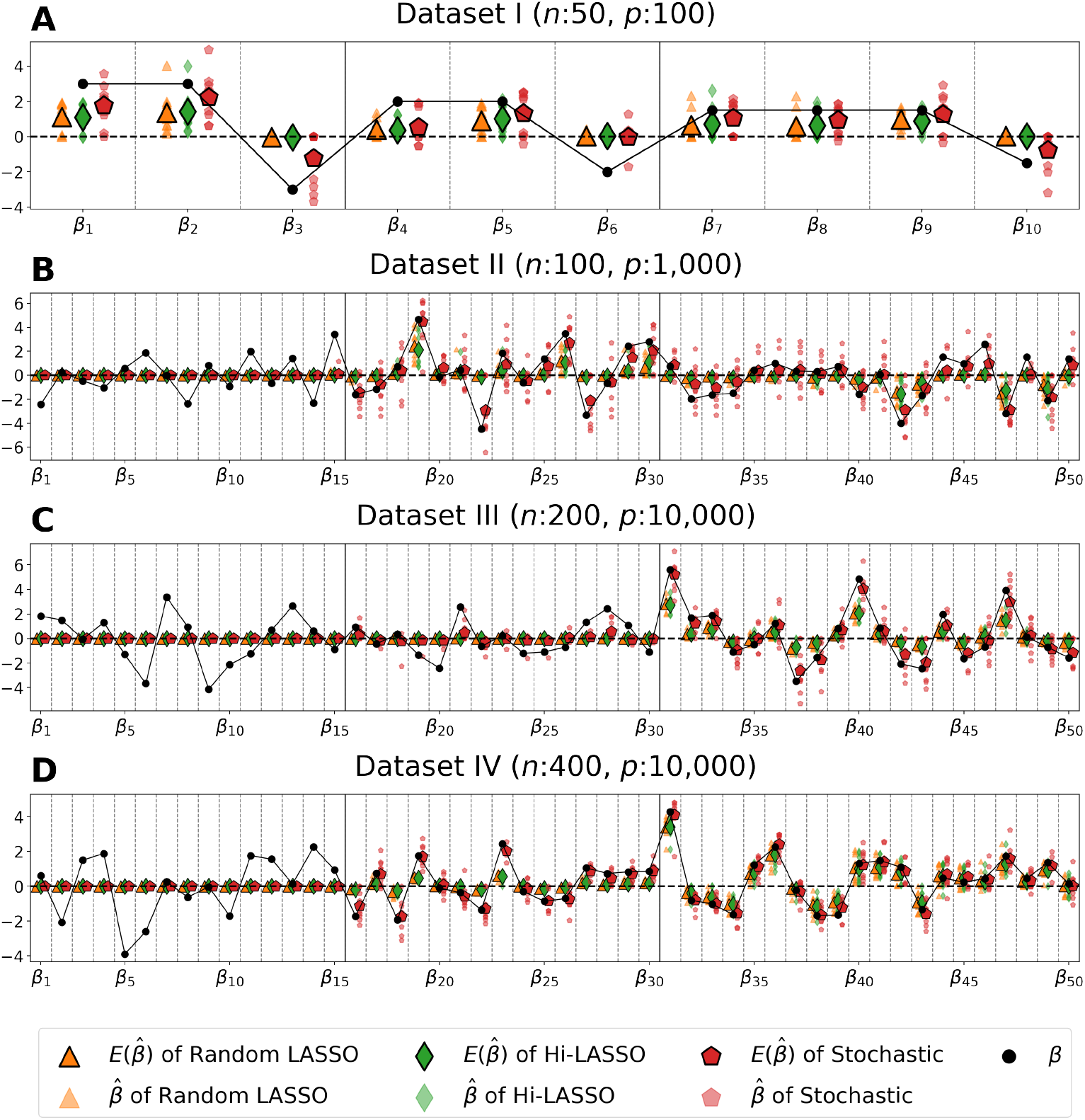
Coefficient estimation with simulation data.

Consequently, the benchmark LASSO models barely identified the negative coefficients (i.e., *β*_3_, *β*_6_, and *β*_10_) in 10 repeated experiments: Random LASSO estimated *β*_3_, *β*_6_, and *β*_10_ as negative coefficients 1, 0, and 0 times, respectively, and Hi-LASSO estimated them 2, 0, and 0 times, respectively. However, Stochastic LASSO demonstrated its notable coefficient estimation capability by accurately estimating *β*_3_, *β*_6_, and *β*_10_ as negative coefficients 4, 2, and 5 times, respectively. We also observed that Random LASSO and Hi-LASSO tend to estimate the same signs in highly collinearity. For instance, In Dataset II, Random LASSO and Hi-LASSO identified only the positive coefficients on *β*_16_ − *β*_30_, and only the negative coefficients on *β*_31_ − *β*_50_, while Stochastic LASSO correctly estimated the signs (Fig. 3B). In Datasets III-IV, only Stochastic LASSO precisely identified *β*_16_−*β*_30_, while Random LASSO and Hi-LASSO underestimated *β*_16_ − *β*_30_ close to zero (Fig. 3C-D).

### 3.3 3.3. Tests of significance

We performed a semi-simulation study to assess whether the proposed tests of significance can identify the statistical significance of non-zero variables in EHDLSS data. In this study, we generated semi-simulation data, adapting gene expressions of 18 types of cancer in the TCGA databases. The gene expression data (i.e., RNA-seq) were directly used as the independent variables, but the dependent variable (e.g., survival month) was synthetically generated as follows: (1) We conducted a correlation analysis between gene expression and the survival months; (2) We selected 100 genes with the highest Pearson correlation coefficient with survival months; (3) The regression coefficients (*β*) of the 100 genes were randomly generated from the normal distribution, *N* (0, 4); (4) The coefficients of the other genes were set to 0; (5) The dependent variables were generated from the linear combination of the gene expression (*X*), the coefficients (*β*), and the errors (*ϵ*) from the normal distribution with a mean of zero and the standard deviation of the logarithmic survival months (i.e., *y* = *X* + *ϵ*). The semi-synthetic cancer datasets are briefly summarized in Table 2. In this experiment, we considered only Recursive Random LASSO and Hi-LASSO, which proposed tests of significance scheme for feature selection, as a benchmark. We computed the F1-scores to evaluate the tests of significance at a significance level of *α* = 0.05.

**Table 2.**
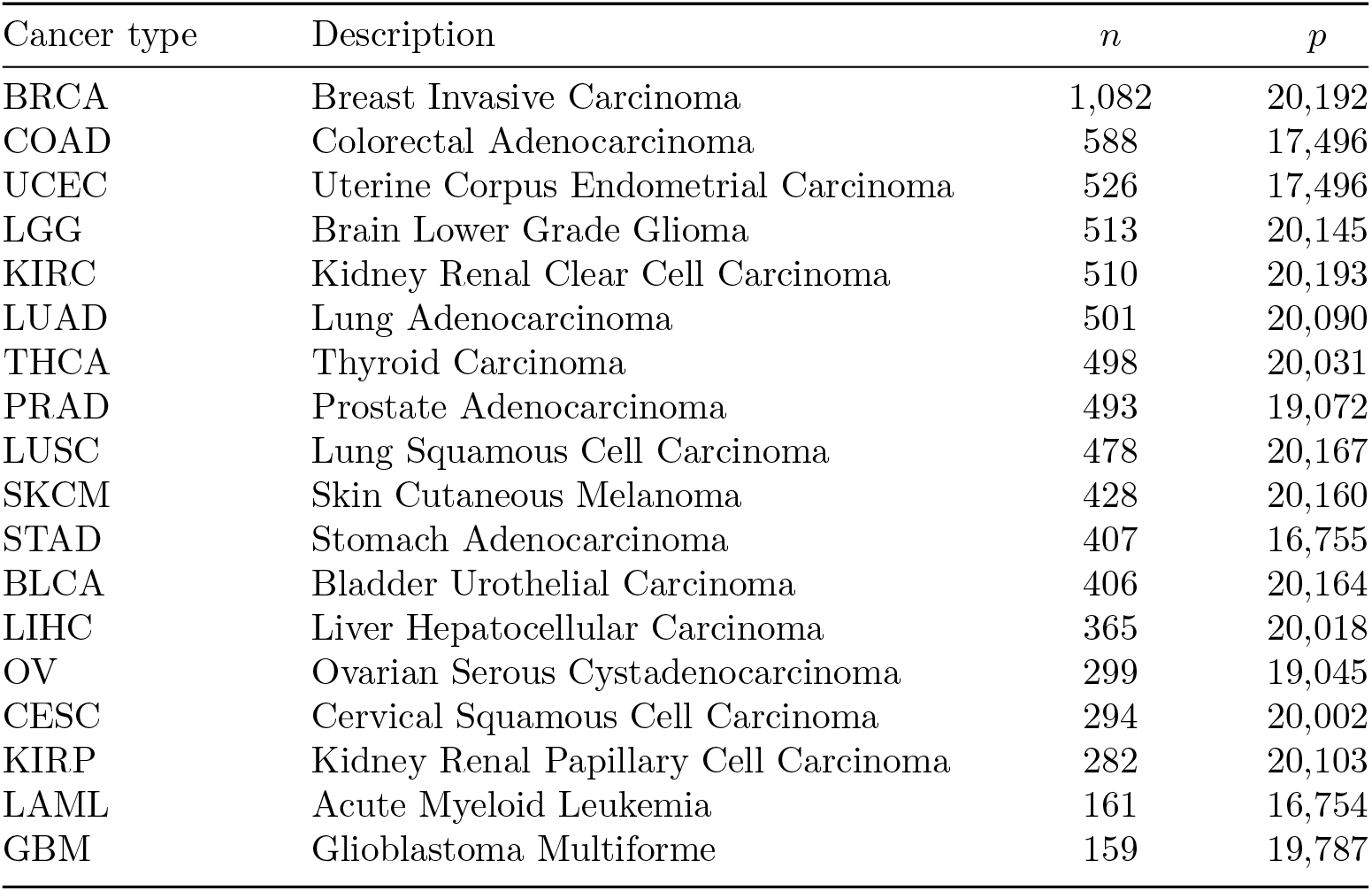
Description of the semi-simulation data.

Stochastic LASSO precisely identified statistically significant non-zero variables, achieving the highest F1-scores across the 18 datasets (Fig. 4, Supplementary Table S3). Stochastic LASSO produced F1-scores of above 0.8 for Breast Invasive Carcinoma (BRCA), Colorectal Adenocarcinoma (COAD), Uterine Corpus Endometrial Carcinoma (UCEC), and Brain Lower Grade Glioma (LGG) datasets, each of which consists of more than 500 samples. Stochastic LASSO also maintained F1-scores of over 0.5 for Glioblastoma Multiforme (GBM) datasets, where the number of samples (*n* : 159) is extremely small compared to the number of features (*p* : 19,787). In contrast, the second-best Hi-LASSO’s performance distinctly declined as the sample sizes decreased. Hi-LASSO showed F1-scores of over 0.5 in the largest sample size datasets only (i.e, BRCA and COAD). These experimental results demonstrate that Stochastic LASSO can provide a reliable feature selection even in extremely high dimensional settings.

**Fig. 4.**
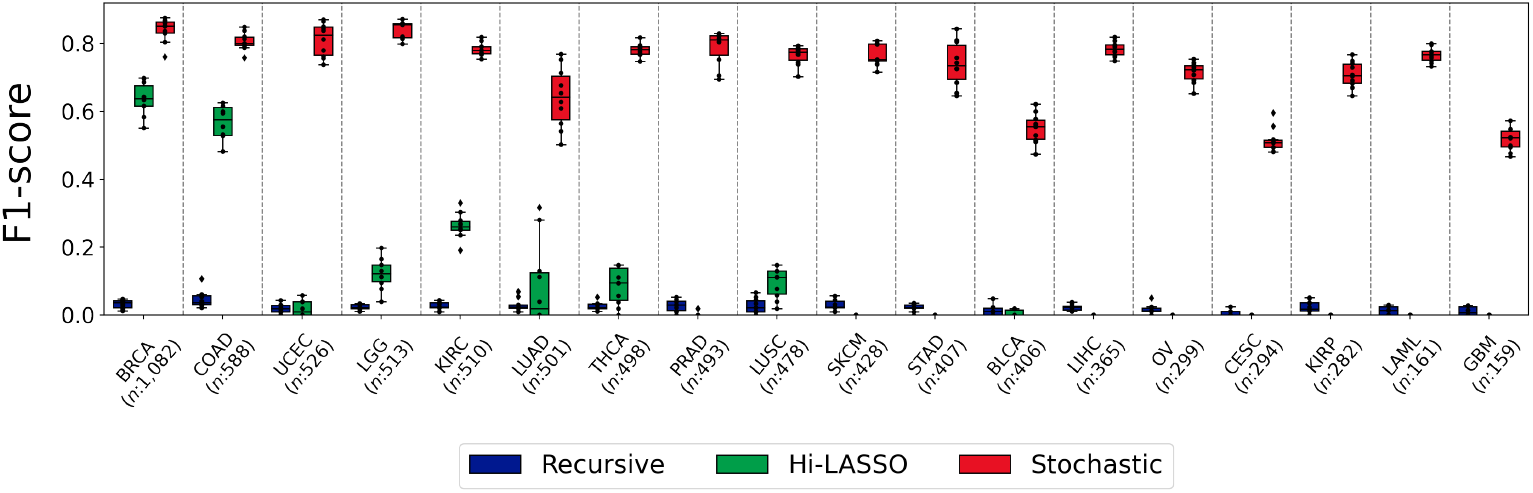
F1-score of the semi-simulation study. The cancer type and sample size (*n*) for each dataset are described on the x-axis of the figure. The number of features (*p*) of each dataset are around 20,000.

### 3.4 3.4. Robustness of feature selection

We finally assessed whether our bootstrap-based LASSO can produce consistent feature selection performance by computing a pair-wise Kuncheva Index (KI) on the previous semi-synthetic cancer datasets. KI computes a robustness score in the range of [-1, 1], where zero indicates that each selection was made independently, a positive value indicates the feature selection model produces a stable selection, and a negative value indicates the model is unstable [17]. The equation of KI is:

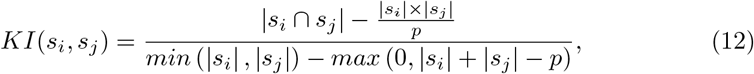

where *s*_*i*_ and *s*_*j*_ are two sets of feature selected by the model, and *p* is the dimensionality of the dataset.

Stochastic LASSO produced the most consistent feature selection with the highest average KI of 0.7590 with smallest variances across 18 cancer types, whereas Hi-LASSO and Recursive Random LASSO showed 0.2158 and 0.0058, respectively (Fig. 5, Supplementary Table S4). Note that Stochastic LASSO achieved the highest F1-scores on the same datasets. The highest F1-scores and KI demonstrate Stochastic LASSO’s reliable feature selection capability for extremely high-dimensional genomic data.

**Fig. 5.**
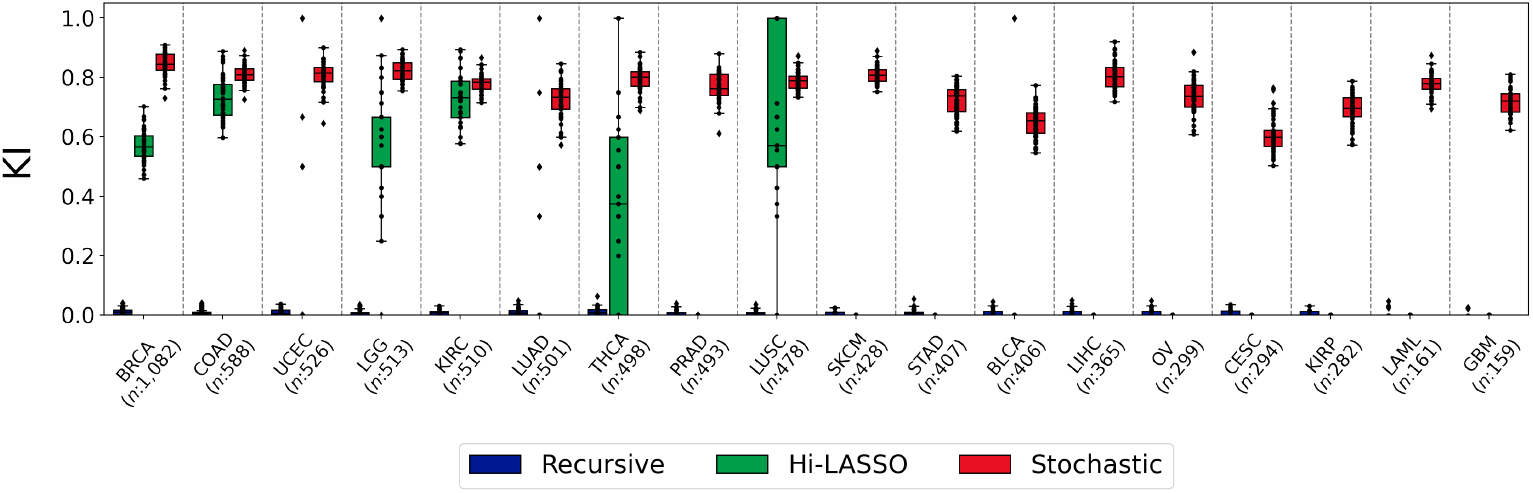
Kuncheva Index (KI) of the semi-simulation study. The cancer type and sample size (*n*) for each dataset are described on the x-axis of the figure. The number of features (*p*) of each dataset is around 20,000.

## 4 Glioblastoma & Glioma Gene Expression Data Analysis

We applied Stochastic LASSO for Glioblastoma Multiforme (GBM) & Brain Lower Grade Glioma (LGG) to assess it with real-world biological data, where ground truths are not known. GBM is the most aggressive and common primary brain tumor in adults, while LGG is a slower-growing, less aggressive brain tumor type with a generally better prognosis [18]. We downloaded genomic data of GBM and LGG from The Cancer Genome Atlas Program (TCGA) and combined them. In the experiment, we used the gene expressions (i.e., RNA-seq) of GBM & LGG patients as independent variables and their survival time as the dependent variable. The gene expression data consist of 266 patients and 19,785 genes, including only deceased patients for the regression problem rather than cox-regression. The average survival time was 28.1 months. We identified genes related to the survival time using Stochastic LASSO, where *q* was set to 266 (i.e., number of sample) and *r* was set to 30 to ensure the normality of coefficient estimates. For the feature selection, we used a significance level of 0.05 for the tests of significance on Stochastic LASSO.

Stochastic LASSO identified 490 statistically significant genes out of 19,785 genes, and we examined the genes in the biological literature. Consequently, a number of genes are shown as associated with biomarkers in GBM & LGG (Table 3). Among the top 20 ranked significant genes identified by Stochastic LASSO, ONECUT3 was reported to suppress glioblastoma cell proliferation and promote a glial-to-neuronal identity switch, implicating it in GBM reprogramming [19]. HILS1 was identified as a strong prognostic biomarker for LGG. It was significantly upregulated in glioma tissues and associated with higher tumor grade and worse survival outcomes [20, 21]. HILS1 was also a part of the five-pseudogene prognostic signatures for lower-grade gliomas, with higher expression levels associated with advanced tumor grade and poorer patient survival [22]. UCN3 was found to be upregulated upon serum stimulation in glioma cells and transiently increased following proliferative stimuli, suggesting a role in glioma adaptation to environmental stress [23, 24]. COL22A1 was overexpressed in GBM and identified as a key angiogenesis-related gene in grade 4 diffuse gliomas. Elevated COL22A1 expression was consistently associated with poor overall survival and endothelial remodeling in the GBM microenvironment [25–27]. LINC00114 was implicated in temozolomide resistance in GBM through ceRNA network regulation [28]. VN1R4 exhibited recurrent genomic alterations in glioma, particularly in LGG, suggesting its association with gliomagenesis and structural genome instability [29].

**Table 3.**
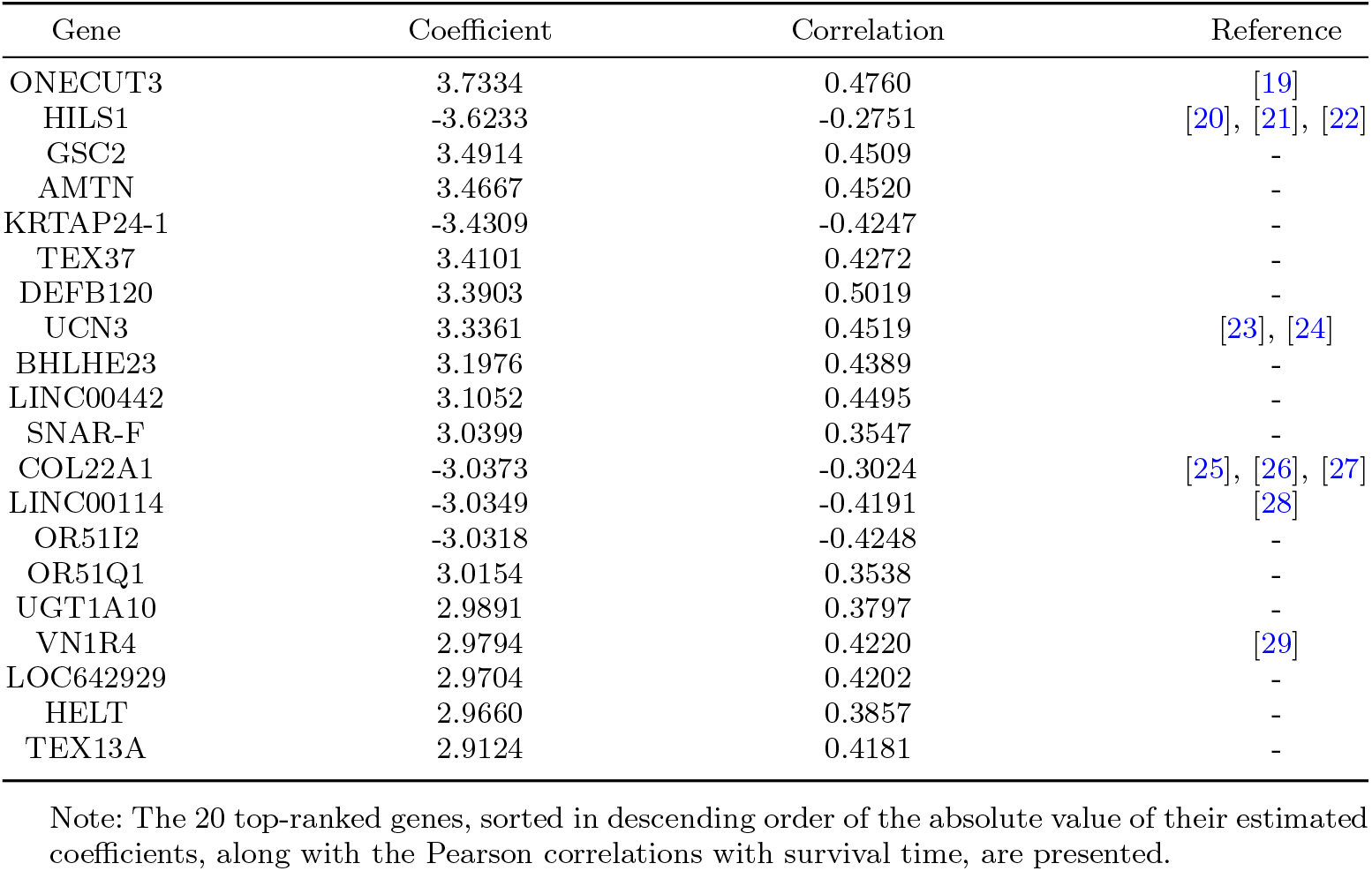
Top-10 ranked genes by Stochastic LASSO in GBM & LGG.

Furthermore, we conducted gene set enrichment analysis (GSEA) using 490 genes identified by Stochastic LASSO, based on 844 curated pathways retrieved from the KEGG database. The statistical significance of enrichment was assessed using the hypergeometric test. Consequently, 13 significantly enriched pathways (p-value*<*0.05) were identified, of which seven pathways have been previously reported to be closely associated with the prognosis of GBM and LGG (Table 4). The olfactory transduction pathway was identified as the most prominently enriched in long-term glioblastoma survivors, as revealed by transcriptomic and epigenetic analyses [30]. Translation initiation was found to be overrepresented in glioblastoma, indicating its relevance to tumor progression and patient outcomes [31]. The ribosome pathway was reported to be significantly downregulated in glioblastoma cells with acquired temozolomide resistance [32]. Retinol metabolism was also downregulated in glioma, suggesting that dysregulation in retinoid processing may contribute to remodeling of the tumor immune microenvironment [33]. The drug metabolism – cytochrome P450 pathway showed notable enrichment in glioma-associated hub genes, indicating a potential link to altered drug response and resistance mechanisms [34]. The metabolism of xenobiotics by cytochrome P450 pathway was enriched in glioblastoma, implicating detoxification processes in therapeutic resistance [35]. The JAK/STAT signaling pathway was reported to play a central role in glioma pathobiology by regulating tumor growth, invasion, immune evasion, and stemness, and thus represented a potential therapeutic target [36].

**Table 4.** Enriched pathway by Stochastic LASSO in GBM & LGG.

| Pathways | Pathway genes | Selected genes | P-values | Reference |
| --- | --- | --- | --- | --- |
| Olfactory Transduction | 389 | 44 | 1.11e-16 | [30] |
| Translation Initiation | 81 | 13 | 9.11e-08 | [31] |
| Ribosome | 88 | 13 | 2.48e-07 | [32] |
| Ascorbate and Aldarate Metabolism | 25 | 5 | 3.22e-04 | - |
| Pentose and Glucuronate Inter-conversions | 28 | 5 | 5.60e-04 | - |
| Retinol Metabolism | 64 | 7 | 1.01e-03 | [33] |
| Drug Metabolism other Enzymes | 51 | 6 | 1.56e-03 | - |
| Drug Metabolism-Cytochrome P450 | 72 | 7 | 2.01e-03 | [34] |
| Steroid Hormone Biosynthesis | 55 | 6 | 2.32e-03 | - |
| Porphyrin and Chlorophyll Metabolism | 41 | 5 | 3.28e-03 | - |
| Metabolism of Xenobiotics by Cytochrome P450 | 70 | 6 | 7.71e-03 | [35] |
| Starch and Sucrose Metabolism | 52 | 5 | 9.12e-03 | - |
| JAK/STAT Signaling pathway | 155 | 8 | 3.94e-02 | [36] |

Note: Significantly enriched pathways identified by Stochastic LASSO (p-value<0.05) are listed, along with the total number of pathway-related genes and the number of genes selected by Stochastic LASSO.

## 5 Conclusions

In this study, we have proposed Stochastic LASSO, an enhanced LASSO model for feature selection with high-throughput data. Stochastic LASSO improves the bootstrap-based LASSO models by: (1) reducing multicollinearity within bootstrap samples while ensuring that each predictor is included an equal number of times, (2) mitigating randomness in predictor sampling by forward selection without additional bootstrapping procedures, and (3) improving statistical significance tests with the proposed two-stage t-test. The performance of Stochastic LASSO was compared to the state-of-the-art LASSO models in extensive simulation settings and with real genomic data. Stochastic LASSO outperformed the benchmarks for feature selection, coefficient estimation, tests of significance, and robustness in the experiments. Stochastic LASSO was applied to gene expression data from TCGA GBM and LGG, identifying both statistically significant genes and enriched pathways associated with survival times. Stochastic LASSO can be applied to any linear regression based models, including survival analysis, protein–protein interactions analysis, and association studies. Furthermore, the application of Stochastic LASSO can be extended to classification analysis by using penalized logistic regression.

## 6 Acknowledgments

This work was supported by the National Science Foundation Major Research Instrumentation (NSF MRI) (Grant#:2117941), the National Research Foundation of Korea (NRF) (NRF-2021R1I1A3048029), and the MSIT (Ministry of Science and ICT) under the ICAN (ICT Challenge and Advanced Network of HRD) support program (IITP-2024-RS-2022-00156409) supervised by the IITP (Institute for Information & Communications Technology Planning & Evaluation) in South Korea.

## 7 Availability of data and code

The datasets analyzed during the current study are all publicly available online from The Cancer Genome Atlas (TCGA). The open-source is available at: https://github.com/datax-lab/StochasticLASSO.

