## Supplementary for "Stochastic LASSO for extremely high-dimensional genomic data"

### Supplementary tables

In every experiment, we underscored the best performing method in bold, and the second best in bold without underlining it. We also conducted a Wilcoxon signed-rank test, which is a non-parametric paired test for the best and second best method (p-value < 0.05 : $*$, p-value < 0.01 : $**$).

**Table S1. Feature selection performance with simulation data.**

| | | **LASSO** | **Elastic** | **Adaptive** | **Relaxed** | **Precision** | **Random** | **Recursive** | **Hi-LASSO** | **Stochastic LASSO** |
|---|---|---|---|---|---|---|---|---|---|---|
| Dataset I | F1 Score | 0.3169<br>(0.0089) | 0.3505<br>(0.0091) | 0.3647<br>(0.0125) | 0.4699<br>(0.0145) | **0.6551**<br>**(0.0134)** | 0.5637<br>(0.0154) | 0.3985<br>(0.0081) | 0.6024<br>(0.0118) | **0.7093**<br>**(0.0197)** |
| | AUCPR | 0.4889<br>(0.0091) | **0.7221**<br>**(0.0111)** | 0.4800<br>(0.0097) | 0.4874<br>(0.0086) | 0.6315<br>(0.0136) | 0.6789<br>(0.0100) | 0.3948<br>(0.0078) | 0.6809<br>(0.0091) | **0.8284****<br>**(0.0095)** |
| Dataset II | F1 Score | 0.1634<br>(0.0062) | 0.3626<br>(0.0139) | 0.1764<br>(0.0052) | 0.1621<br>(0.0051) | 0.2111<br>(0.0193) | 0.3408<br>(0.0110) | 0.0672<br>(0.0034) | **0.3761**<br>**(0.0117)** | **0.6550****<br>**(0.0139)** |
| | AUCPR | 0.1709<br>(0.0036) | 0.3910<br>(0.0126) | 0.1660<br>(0.0031) | 0.1630<br>(0.0023) | 0.4490<br>(0.0081) | 0.4983<br>(0.0071) | 0.0948<br>(0.0025) | **0.5098**<br>**(0.0064)** | **0.6992****<br>**(0.0021)** |
| Dataset III | F1 Score | 0.0902<br>(0.0043) | 0.1831<br>(0.0157) | 0.1281<br>(0.0031) | 0.0916<br>(0.0057) | 0.0602<br>(0.0060) | **0.3175**<br>**(0.0138)** | 0.0264<br>(0.0026) | 0.2749<br>(0.0151) | **0.5251****<br>**(0.0169)** |
| | AUCPR | 0.0726<br>(0.0020) | 0.3498<br>(0.0073) | 0.1014<br>(0.0047) | 0.0674<br>(0.0027) | 0.3413<br>(0.0169) | **0.4389**<br>**(0.0046)** | 0.0198<br>(0.0014) | 0.4354<br>(0.0048) | **0.5772****<br>**(0.0089)** |
| Dataset IV | F1 Score | 0.2155<br>(0.0079) | 0.2506<br>(0.0145) | 0.2709<br>(0.0040) | 0.2495<br>(0.0069) | 0.2380<br>(0.0264) | 0.4776<br>(0.0170) | 0.0420<br>(0.0028) | **0.5482**<br>**(0.0161)** | **0.7777****<br>**(0.0046)** |
| | AUCPR | 0.1757<br>(0.0032) | 0.5083<br>(0.0052) | 0.2155<br>(0.0048) | 0.1649<br>(0.0030) | 0.6174<br>(0.0051) | 0.6434<br>(0.0027) | 0.0277<br>(0.0015) | **0.6492**<br>**(0.0028)** | **0.6985****<br>**(0.0018)** |

**Table S2. Coefficient estimation performance with simulation data.**

| | | **LASSO** | **Elastic** | **Adaptive** | **Relaxed** | **Precision** | **Random** | **Recursive** | **Hi-LASSO** | **Stochastic LASSO** |
|---|---|---|---|---|---|---|---|---|---|---|
| Dataset I | $RMSE_{ALL}$ | 0.5925<br>(0.0041) | 0.5943<br>(0.0036) | 0.5988<br>(0.0035) | 0.5899<br>(0.0047) | 0.5685<br>(0.0039) | 0.5622<br>(0.0034) | 0.6558<br>(0.0111) | **0.5549**<br>**(0.0032)** | **0.5022***<br>**(0.0091)** |
| | $RMSE_{Nonzeros}$ | 1.8113<br>(0.0147) | 1.8349<br>(0.0120) | 1.8145<br>(0.0129) | 1.8325<br>(0.0155) | 1.7787<br>(0.0125) | 1.7664<br>(0.0108) | 2.0641<br>(0.0348) | **1.7450**<br>**(0.0097)** | **1.5286***<br>**(0.0294)** |
| Dataset II | $RMSE_{ALL}$ | 0.3995<br>(0.0009) | 0.4154<br>(0.0003) | 0.3993<br>(0.0011) | 0.3968<br>(0.0009) | 0.3884<br>(0.0022) | 0.3819<br>(0.0007) | 0.4587<br>(0.0027) | **0.3801**<br>**(0.0008)** | **0.3596**<br>**(0.0041)** |
| | $RMSE_{Nonzeros}$ | 1.7718<br>(0.0049) | 1.8412<br>(0.0014) | 1.7749<br>(0.0054) | 1.7597<br>(0.0038) | 1.7258<br>(0.0104) | 1.7063<br>(0.0033) | 2.0365<br>(0.0125) | **1.6976**<br>**(0.0035)** | **1.5931**<br>**(0.0179)** |
| Dataset III | $RMSE_{ALL}$ | 0.1294<br>(0.0002) | 0.1405<br>(0.0002) | 0.1265<br>(0.0003) | 0.1298<br>(0.0003) | 0.1331<br>(0.0001) | **0.1207**<br>**(0.0002)** | 0.1510<br>(0.0007) | 0.1211<br>(0.0003) | **0.1099****<br>**(0.0006)** |
| | $RMSE_{Nonzeros}$ | 1.8224<br>(0.0025) | 1.9788<br>(0.0025) | 1.7830<br>(0.0045) | 1.8247<br>(0.0030) | 1.8808<br>(0.0019) | **1.7067**<br>**(0.0028)** | 2.1177<br>(0.0105) | 1.7128<br>(0.0039) | **1.5464****<br>**(0.0098)** |
| | $RMSE_{ALL}$ | 0.0943<br>(0.0001) | 0.1029<br>(0.0001) | 0.0924<br>(0.0002) | 0.0935<br>(0.0001) | 0.0976<br>(0.0002) | **0.0841**<br>**(0.0001)** | 0.1231<br>(0.0012) | 0.0842<br>(0.0001) | **0.0777****<br>**(0.0002)** |

| | | | | | | | | | |
|---|---|---|---|---|---|---|---|---|---|
| Dataset IV | $RMSE_{Nonzeros}$ | 1.3315 (0.0008) | 1.4516 (0.0009) | 1.3038 (0.0023) | 1.3214 (0.0014) | 1.3772 (0.0033) | **1.1882 (0.0017)** | 1.7363 (0.0180) | 1.1907 (0.0021) | **1.0945** (0.0025)** |

**Table S3. F1-score with semi-simulation data.**

| Cancer type | $n$ | $p$ | Recursive | Hi-LASSO | Stochastic LASSO |
|---|---|---|---|---|---|
| BRCA | 1,082 | 20,192 | 0.0327 (0.0013) | **0.6363 (0.0048)** | **0.8398 (0.0035)**** |
| COAD | 588 | 17,496 | 0.0474 (0.0025) | **0.5639 (0.0056)** | **0.8057 (0.0026)**** |
| UCEC | 526 | 17,496 | 0.0200 (0.0013) | **0.0215 (0.0025)** | **0.8112 (0.0049)**** |
| LGG | 513 | 20,145 | 0.0236 (0.0009) | **0.1228 (0.0046)** | **0.8405 (0.0026)**** |
| KIRC | 510 | 20,193 | 0.0271 (0.0012) | **0.2632 (0.0038)** | **0.7827 (0.0020)**** |
| LUAD | 501 | 20,090 | 0.0291 (0.0019) | **0.0880 (0.0122)** | **0.6409 (0.0089)**** |
| THCA | 498 | 20,031 | 0.0259 (0.0013) | **0.0859 (0.0055)** | **0.7806 (0.0019)**** |
| PRAD | 493 | 19,072 | **0.0283 (0.0017)** | 0.0040 (0.0008) | **0.7874 (0.0051)**** |
| LUSC | 478 | 20,167 | 0.0275 (0.0022) | **0.0972 (0.0046)** | **0.7654 (0.0029)**** |
| SKCM | 428 | 20,160 | **0.0302 (0.0017)** | 0.0000 (0.0000) | **0.7658 (0.0032)**** |
| STAD | 407 | 16,755 | **0.0220 (0.0011)** | 0.0000 (0.0000) | **0.7393 (0.0066)**** |
| BLCA | 406 | 20,164 | 0.0155 (0.0019) | **0.0059 (0.0010)** | **0.5499 (0.0044)**** |
| LIHC | 365 | 20,018 | **0.0227 (0.0009)** | 0.0000 (0.0000) | **0.7832 (0.0022)**** |
| OV | 299 | 19,045 | **0.0160 (0.0014)** | 0.0000 (0.0000) | **0.7141 (0.0031)**** |
| CESC | 294 | 20,002 | **0.0073 (0.0010)** | 0.0000 (0.0000) | **0.5148 (0.0036)**** |
| KIRP | 282 | 20,103 | **0.0234 (0.0019)** | 0.0000 (0.0000) | **0.7083 (0.0038)**** |
| LAML | 161 | 16,754 | **0.0131 (0.0013)** | 0.0000 (0.0000) | **0.7662 (0.0022)**** |
| GBM | 159 | 19,787 | **0.0115 (0.0013)** | 0.0000 (0.0000) | **0.5175 (0.0033)**** |

**Table S4. KI with semi-simulation data.**

| Cancer type | $n$ | $p$ | Recursive | Hi-LASSO | Stochastic LASSO |
|---|---|---|---|---|---|
| BRCA | 1,082 | 20,192 | 0.0108 (0.0011) | **0.5698 (0.0054)** | **0.8453 (0.0038)**** |
| COAD | 588 | 17,496 | 0.0091 (0.0012) | **0.7321 (0.0068)** | **0.8082 (0.0032)**** |
| UCEC | 526 | 17,496 | 0.0077 (0.0012) | **0.0814 (0.0242)** | **0.8081 (0.0047)**** |
| LGG | 513 | 20,145 | 0.0076 (0.0013) | **0.5609 (0.0208)** | **0.8222 (0.0037)**** |
| KIRC | 510 | 20,193 | 0.0071 (0.0011) | **0.7227 (0.0083)** | **0.7805 (0.0032)**** |
| LUAD | 501 | 20,090 | 0.0077 (0.0013) | **0.1313 (0.0269)** | **0.7267 (0.0063)**** |
| THCA | 498 | 20,031 | 0.0088 (0.0015) | **0.4045 (0.0316)** | **0.7943 (0.0042)**** |
| PRAD | 493 | 19,072 | **0.0035 (0.0011)** | 0.0000 (0.0000) | **0.7678 (0.0049)**** |
| LUSC | 478 | 20,167 | 0.0049 (0.0011) | **0.6157 (0.0306)** | **0.7889 (0.0031)**** |

| | | | | | |
|---|---|---|---|---|---|
| SKCM | 428 | 20,160 | **0.0033 (0.0009)** | 0.0000 (0.0000) | **0.8085 (0.0028)**** |
| STAD | 407 | 16,755 | **0.0071 (0.0013)** | 0.0000 (0.0000) | **0.7220 (0.0048)**** |
| BLCA | 406 | 20,164 | 0.0053 (0.0012) | **0.0667 (0.0249)** | **0.6485 (0.0049)**** |
| LIHC | 365 | 20,018 | **0.0065 (0.0013)** | 0.0000 (0.0000) | **0.8077 (0.0046)**** |
| OV | 299 | 19,045 | **0.0049 (0.0013)** | 0.0000 (0.0000) | **0.7368 (0.0052)**** |
| CESC | 294 | 20,002 | **0.0047 (0.0011)** | 0.0000 (0.0000) | **0.6056 (0.0058)**** |
| KIRP | 282 | 20,103 | **0.0022 (0.0010)** | 0.0000 (0.0000) | **0.6928 (0.0050)**** |
| LAML | 161 | 16,754 | **0.0015 (0.0011)** | 0.0000 (0.0000) | **0.7772 (0.0037)**** |
| GBM | 159 | 19,787 | **0.0008 (0.0008)** | 0.0000 (0.0000) | **0.7200 (0.0045)**** |
